# Benchmarking single-cell foundation models for real-world RNA-seq data integration

**DOI:** 10.64898/2026.04.17.719314

**Authors:** Siyu Han, Tamas Sztanka-Toth, Enes Senel, Ahmed Elnaggar, Jaymala Patel, Tommaso Mansi, Denis Smirnov, Joel Greshock, Alex Javidi

## Abstract

Single-cell foundation models enable reusable representations and streamlined analysis workflows, yet rigorous evaluation of their performance and robustness in real-world pharmaceutical settings remain underexplored. Here, we benchmarked leading single-cell foundation models (scGPT; scGPT_CP, a continually pretrained checkpoint of scGPT; scFoundation; scMulan; CellFM) against established baseline methods (scVI; Harmony) for data integration using over 1.5 million cells from clinical and preclinical samples. Performance was assessed using well-established and complementary metrics for technical correction and biological structure preservation. We further introduced robustness-oriented rankings to summarize metric trade-offs and quantify performance consistency across datasets and evaluation settings. Our findings show that fine-tuning improved technical correction performance; among the foundation models, fine-tuned scGPT_CP performed best. However, the baseline scVI was the top overall performer, ranking first by our multi-metric Leximax ranking and achieving the highest Pareto Front-1 hit. Collectively, our study provides practical insights for adapting foundation models to real-world drug design and development.

## Introduction

Single-cell transcriptomics is widely used across diverse biomedical applications, including drug discovery and development^1,2^. Real-world datasets spanning donors, batches, and assays involve technical variation that can mask true biological signals. Reliable data integration is therefore critical for robust downstream analyses and well-grounded clinical decisions. Despite advances in single-cell research, effective data integration remains an open problem highlighted by recent community benchmarks^3–5^.

Single-cell foundation models (scFMs) have been proposed as a promising approach to address this challenge by providing reusable cell embeddings for diverse downstream tasks. Leveraging curated public datasets, recent benchmarking efforts have evaluated scFM performance on data integration and revealed both scFM benefits and limitations^6,7^. Since atlas-style public resources used in these studies cannot fully reflect a real-world drug development setting, we complement this field by conducting evaluation on 1.5 million cells from three non-public datasets collected from clinical and preclinical drug development workflows.

Our benchmark datasets include apheresis samples from a diversity of sources. These are i.) multiple myeloma patients treated with CAR-T therapy (Apheresis dataset), ii.) primary and metastatic prostate cancer biopsies and necropsies (Prostate cancer dataset), and iii.) CAR-T drug product (CAR-T drug product dataset) (Methods, Extended Data Table 1). The Apheresis and CAR-T drug product datasets are from standard single-cell RNA sequencing (scRNA), while the Prostate cancer dataset uniquely contains scRNA and single-nucleus (snRNA) data. Clinical research often includes frozen or hard-to-dissociate tissues that are profiled by snRNA. The Prostate cancer dataset is designed to evaluate method’s applicability for this specific scenario. CAR-T manufacturing can alter expression profiles and lead to a data distribution that differs from typical pretraining data used by scFMs. We built the CAR-T drug product dataset to test whether integration remains reliable under this out-of-distribution scenario. Together, these datasets let us evaluate scFMs in settings that better reflect clinical development and cell therapy manufacturing.

We evaluated the widely used scGPT^8^ and scFoundation^9^, as well as three models not assessed in earlier benchmarks: scMulan^10^, CellFM^11^, and scGPT_CP (a continually pretrained scGPT checkpoint released by the original authors)^8^. Where supported, we assessed both zero-shot (use pre-trained model directly without further training) and fine-tuned (with additional parameter updates on benchmark data) configurations to quantify out-of-the-box generalization and data-specific adaptation (Methods). Embeddings from principal component analysis (PCA) and task-specific methods scVI^12^ and Harmony^13^ served as baselines (Methods, Extended Data Table 2). In total, 10 distinct embeddings were included in this study.

We performed sample-level integration and evaluated performance using three complementary metrics: batch correction and bio conservation scores from scib-metrics^14^, and scGraph score^7^ (Methods). Both the batch correction score and the bio conservation score are aggregated metrics, with the former quantifying how well batch-specific variation is removed, while the latter measures whether biological signal is preserved after integration. Existing benchmarks have largely focused on these two metrics alone, potentially overlooking the topology of cell-cell relationships^7^. The recently proposed scGraph score assesses this biological relevance and reveals if the cell-cell distances have been distorted after integration. A high value for these metrics indicates better integration. We used uniform manifold approximation and projection^15^ (UMAP) to visualize cross-assay integration (Methods).

## Results

In our zero-shot analysis (Figure 1a, Supplementary Data Table 1), embeddings from scFMs generally achieved higher batch correction scores than unintegrated PCA embeddings. However, no single scFM consistently ranked best across metrics and datasets. scGPT_CP achieved the highest batch correction scores on the Prostate cancer and CAR-T drug product datasets, whereas scFoundation led on the Apheresis dataset. For bio conservation, scFoundation achieved the highest score on the Apheresis dataset, but scGPT (0.437) outperformed scFoundation (0.414) on the Prostate cancer dataset. On the CAR-T drug product dataset, all scFMs showed lower bio conservation scores than unintegrated PCA. Among these, scFoundation’s bio conservation score was closest to PCA, with a 0.003 decrease, while scMulan showed the largest reduction (0.125). For scGraph scores, scMulan achieved the highest value on the Apheresis dataset (0.772), and scFoundation was second (0.738) on this dataset and led on the other two datasets. Across the datasets and metrics assessed, scFoundation showed stable, strong zero-shot performance, although it was not always the top method for a given metric or dataset. Interestingly, scFoundation improves data quality by mitigating sequencing depth variation and does not have a module tailored for data integration (Methods).

**Figure 1.**
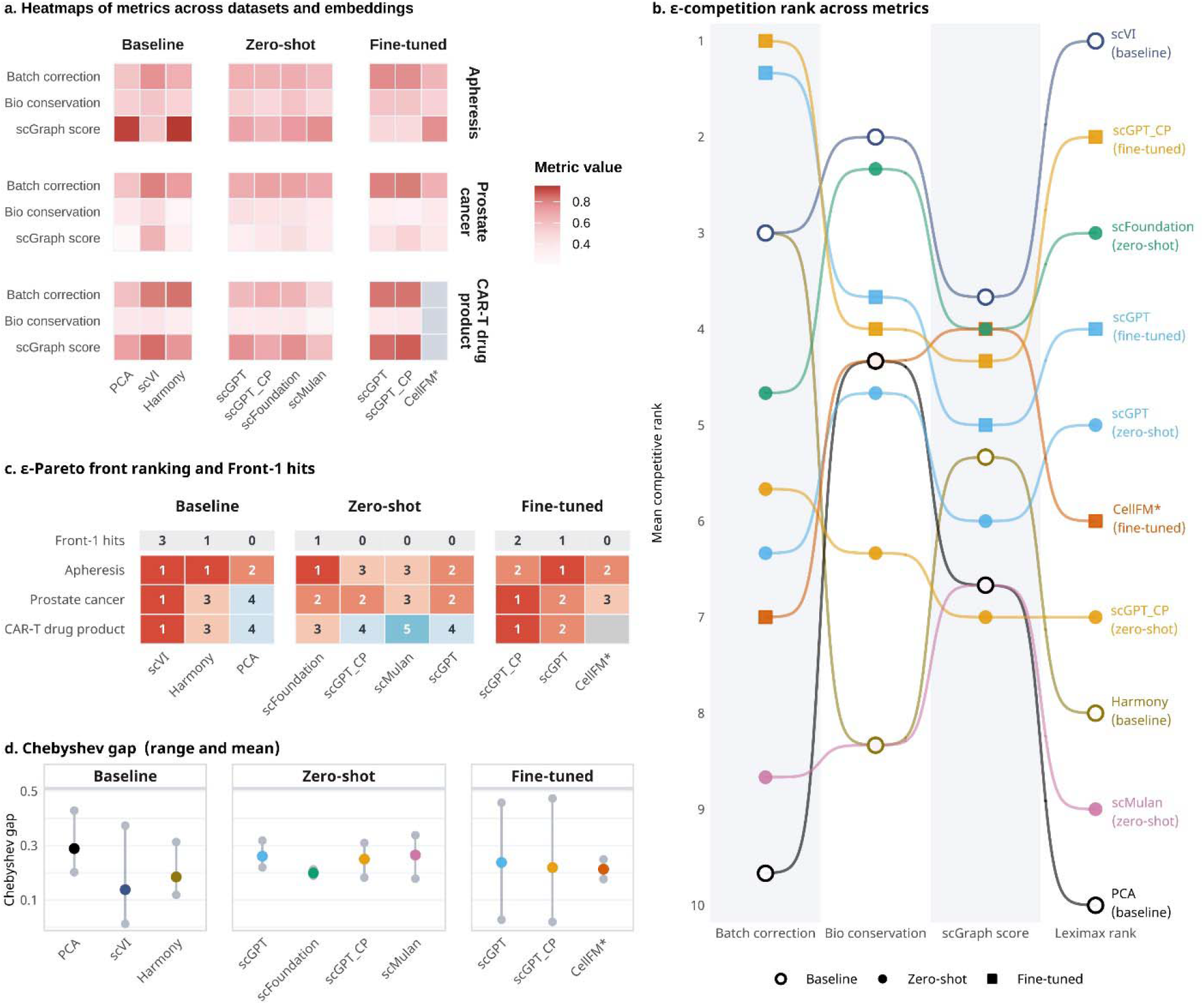
**a**. Heatmap of the metrics values for 10 embeddings across three metrics and three benchmarking datasets. A higher value indicates better integration results. **b**, Bump chart of mean competition ranks across metric (baseline methods: PCA, scVI, Harmony; zero-shot scFMs: scGPT, scGPT_CP, scFoundation, scMulan; fine-tuned scFMs: scGPT, scGPT_CP, CellFM). Lower ranks indicate better performance. The mean ranks for batch correction, bio conservation, and scGraph metrics are computed across the benchmarking datasets. The Leximax rank prioritizes methods with the best worst-performing metric. Lower Leximax ranks indicate more balanced performance. **c**, Pareto front ranking of each embedding per dataset. A method in an earlier front dominates a method in a later front if it is not worse on any metric and strictly better on at least one metric. Methods within the same front are mutually non-dominated. A method with more Front-1 hits is more robust in achieving optimal trade-offs among metrics across datasets. **d**, Range plot showing the Chebyshev gap across different methods. Each colored dot represents the mean value, with vertical lines indicating the range. The ranges capture the variability of each method’s performance across

Fine-tuning consistently improved batch correction scores. On the CAR-T drug product dataset, fine-tuned scGPT models outperformed their zero-shot counterparts across all metrics, which suggests fine-tuning is beneficial for data-specific adaptation. We found that scVI tends to better preserve biological information. It achieved the highest bio conservation scores on both the Prostate cancer and CAR-T drug product datasets. On the Apheresis dataset, scVI ranked fourth on this score (0.574), slightly outperformed by fine-tuned scGPT (0.589), zero-shot scFoundation (0.583), and fine-tuned scGPT-CP (0.582). For CellFM, its fine-tuning on the CAR-T drug product exceeded time limits and was excluded. It showed weaker batch correction than scGPT and scGPT_CP on the remaining two datasets, and its biological signal preservation varied across datasets.

As shown in Figure 1a, different methods exhibit variation in performance across datasets and metrics. Given the heterogeneity of datasets and distinct aspects captured by each metric, we argue that a rank-based strategy offers a more robust comparison than absolute metric values to enable decision making about the most suitable scFM under different data contexts and metric priorities. We therefore adopt the following ranking methodology to evaluate both overall performance and cross-metric balance (Supplementary Data Table 2):

- ε-competition ranking: We calculated the average per-metric ranking across all datasets. Methods with metric values differing by less than ε = 0.01 are assigned the same rank to adjust for negligible performance gaps. Building on the concept of lexicographic minimax^16^, we used Leximax rank to summarize balanced performance. Lower Leximax ranks indicate that methods are stable across all metrics, without favoring one metric at the expense of others.
- ε-Pareto front ranking: Methods belonging to lower-ranked fronts indicate better performance. The Front-1 hits show the number of times each method appears on Front 1 across the benchmarking datasets. We also applied a tolerance ε = 0.01 to ignore small performance differences (Fig 1c).
- Chebyshev gap: For each dataset, this measures each method’s worst per-metric gap to the dataset-specific best. We calculated the mean gap and the range across datasets to reveal metric bias and cross-dataset stability (Fig 1d).

Fig 1b shows baseline methods and scFMs (zero-shot and fine-tuned) rankings across metrics, and how these rankings vary between metrics. scGPT_CP (fine-tuned) obtained the highest performance in batch correction. The baseline scVI ranked third in batch correction but demonstrated the strongest performance on bio conservation and scGraph score. scVI also ranked first by the Leximax criterion, which suggests it had the most balanced performance across metrics. By Leximax, scFoundation ranked third overall and first among the zero-shot models. scVI was on Front 1 in all three datasets, showing its robustness across the benchmarking datasets (Fig 1c); the fined-tuned scGPT_CP model was on Front 1 in the Prostate cancer and CAR-T drug product datasets. scVI retained its lead in the Chebyshev gap metric but had relatively larger gap range compared to Harmony and scFM zero-shot models (Fig 1d). scFoundation, in contrast, shows the smallest gap range, indicating its high stability across data.

We finally visualize cross-assay integration using UMAP embeddings, colored by transcriptomic assay and cell type (Fig 2). If integration works well, cells of the same type should largely mix in the embedding space, rather than separating by assay. The results show that all scFMs struggled to integrate the scRNA and snRNA modalities, with cells from the two assays failing to mix properly (Fig 2a). The cell type visualization shows that epithelial and beta cells are largely from snRNA samples (Fig 2b). However, cells of these two populations formed numerous fragments in the embeddings generated by scMulan and CellFM, which suggests that these embeddings were less coherent even within the same modality. By contrast, several other methods, such as Harmony and scVI, while still showing incomplete cross-assay mixing, produced more contiguous clusters within each modality, indicating better within-assay integration. Note that scVI’s cross-assay performance here may be conservative. The assay label can be included as a covariate in scVI. Because scFMs typically do not support additional covariates, we did not provide this information to any method to ensure a fair comparison.

**Figure 2.**
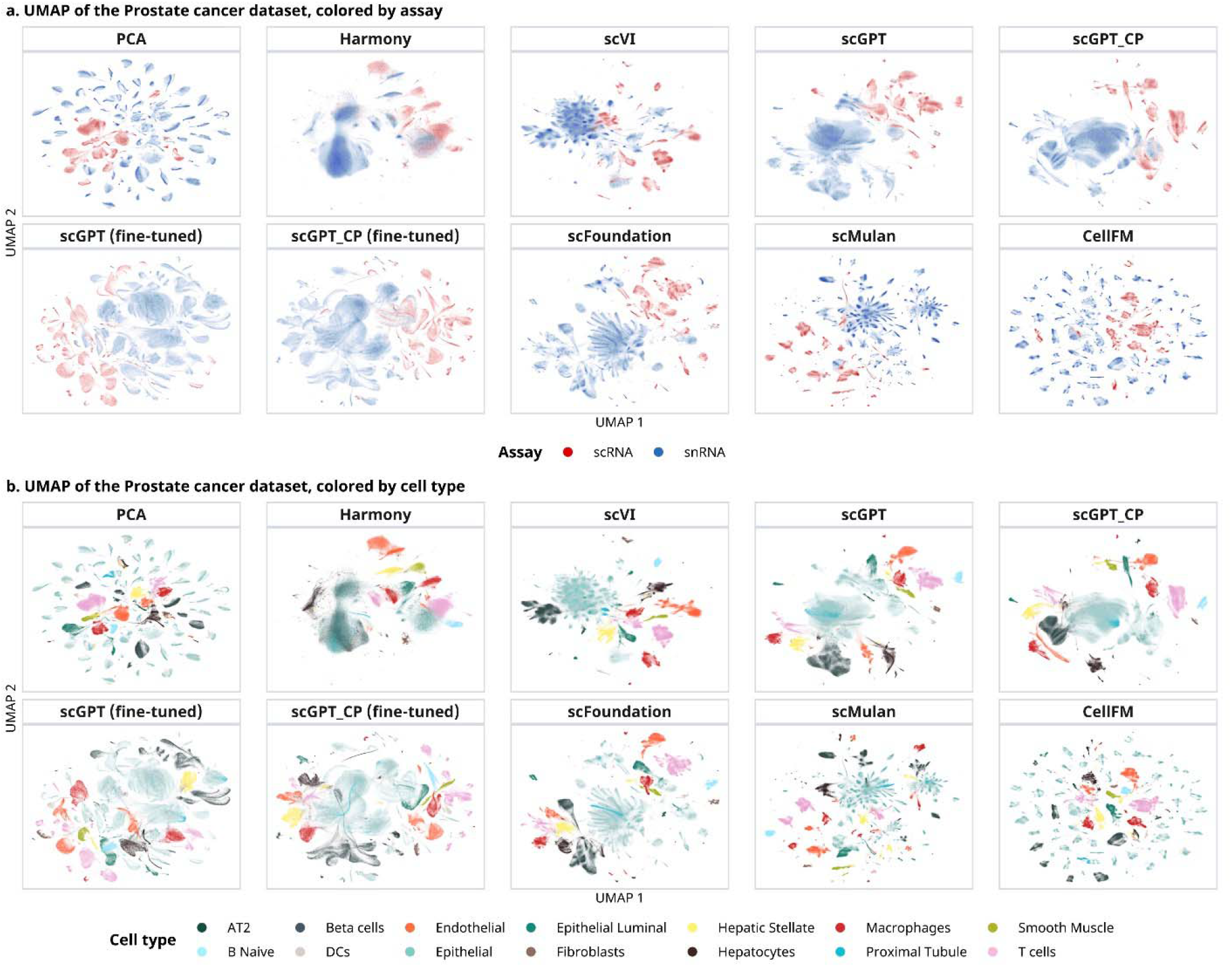
UMAP visualization of cell embeddings from the Prostate cancer dataset, colored by transcriptomic assay (**a**) and cell type (**b**). UMAPs are computed from each embedding separately. Effective integration would mix cells of the same type cross assays, rather than forming assay-separated clusters. Here, all foundation models show limited cross-assay integration (**a**). In the cell type views (**b**), several methods, such as Harmony and scVI, show good within-assay alignment, with cells of the same type clustering together within individual assays.

## Conclusion

Our multi-metric comparison concurs with recent benchmarks^4,17^, showing that even after fine-tuning, current scFMs do not match state-of-the-art task-specific methods like scVI on data integration. When accounting for computational cost (Extend Data Table 3), scFMs do not show advantages for this single task. We note that fine-tuned scGPT_CP and scFoundation did not lag far behind scVI. If embedding can be reused across diverse downstream tasks, scFMs still have great potential to reduce efforts for tuning numerous task-specific models.

Results from scGPT show that fine-tuning improves model capabilities. Given scFoundation’s strong zero-shot performance, its results may further improve once fine-tuning is supported. Our benchmark also reveals scFM’s limited effectiveness for cross-assay integration. There are already task-specific methods, such as sysVI^18,19^, that can be used to integrate datasets with strong technical variations. Future scFMs may benefit from explicit context-aware conditioning for better cross-domain integration.

scFMs have been rapidly expanding our toolbox for biomedical data modeling. Although this benchmark on data integration does not necessarily generalize to other tasks, our findings highlight the importance of aligning these emerging capabilities with real-world needs. In practice, scVI remains the safest default under diverse datasets and evaluation criteria. When embedding reuse is a priority, scFoundation (zero-shot stability; non-commercial research use) and fine-tuned scGPT_CP (stronger data integration; commercial use permitted) can be considered, while performance on the intended downstream tasks should also be verified.

## Methods

### Benchmark datasets

We benchmarked data integration methods on three internal datasets in Johnson & Johnson that are used for real-world drug discovery and development (Extended Data Table 1). The Apheresis dataset contains single-cells from apheresis material of patients receiving CAR-T therapy. The Prostate cancer dataset contains single-cells biopsies and necropsies from primary and metastatic tumors, from prostate cancer patients. Finally, the CAR-T drug product dataset contains single-cells from CAR-T drug product samples, both from healthy donors and patients receiving CAR-T therapy.

To mitigate both technical variation and biological heterogeneity across the patient population, we perform data integration at sample level, treating sample ID as the batch variable. This approach ensures that the learned embeddings have shared biological context for downstream analysis. The cell types of the Prostate cancer dataset were annotated using marker genes, while the Apheresis and CAR-T drug product datasets were annotated using CellTypist^20^ and further validated using marker genes.

In the Prostate cancer dataset, samples include both scRNA-seq and snRNA-seq. Since not all methods accept additional covariates, we did not include assay labels as an explicit covariate to ensure all methods were given the same information. We assume that sample-level differences are fine-grained variation, where the assay-level variation is a coarser, systematic shift. Assay-level shift is expected to be handled implicitly through sample-level integration.

### Software versions and parameters

Where possible, we fine-tuned the models with a maximum of 10 epochs. Early stopping was enabled when supported by the original implementation. All benchmarked methods were evaluated using the hyperparameters suggested in their documentation:

- scGPT supports both zero-shot and fine-tuning integration. The script version for scGPT is 0.2.4. Both the original human model (scGPT_human, released 2023-06-23) and continually pretrained model (scGPT_CP, reseased 2023-12-19) were evaluated. We followed the tutorial for zero-shot (<u>tutorial for zero-shot integration</u>) and fine-tuned integration (<u>tutorial for fine-tuned integration</u>).
- scFoundation (main branch, model released February 2025) only supports zero-shot learning and does not provide a built-in data integration function. In our benchmark, we applied the default read depth enhancement module (target resolution = 4) as described in the original publication to harmonize the cell with different read depths. We aimed to evaluate the extent to which this approach could mitigate technical variation within the datasets.
- scMulan currently supports only zero-shot integration. We used the latest scripts and parameter settings (https://github.com/cyxss/scMulan, Commit d4ddbfb, 2024-11-23) for data integration.
- For CellFM (main branch, MindSpore implementation), data integration was performed using the CellFM-80M model (December 17, 2024) following the procedures outlined in the tutorial (<u>CellFM/tutorials/BatchIntegration/BatchIntegration.ipynb at main · biomed-AI/CellFM</u>).

Two task-specific methods were used as baselines:

- scVI (v1.3.2) was used as a baseline method, with two hidden layers a latent space dimensionality of 30, as recommended in the tutorial (<u>tutorial</u>). In accordance with established practices, 2,000 highly variable genes were selected for integration^14^.
- Harmony (implementation in Scanpy^21^ v1.10.4), a self-supervised data integration method based on embeddings was employed with default settings. Principle components were provided as input.

### Evaluation metrics

We evaluated integrated cell embeddings using metrics from scib-metrics and the scGraph score. Scib-metrics emphasizes batch mixing and within-cell-type information preservation, while scGraph score evaluates cross-cell-type topology. These metrics capture complementary aspects and can be categorized into three groups for a complete assessment of integration quality.

#### Batch correction metrics

We use Average Silhouette Width by batch (ASW_batch), Graph Connectivity (GC), and principal component regression (PCR) comparison to evaluate how well cells from different batches mix together. ASW_batch measures mixing of batches, while GC evaluates whether cells of the same type remain connected across batches in the *k*-NN graph. PCR comparison is used to compare the explained variance before and after integration. The aggregated batch correction score is computed as the arithmetic mean across all three batch correction metrics.

#### Bio-conservation metrics

We use Normalized mutual information (NMI), Adjusted Rand Index (ARI) and Average Sihouette Width by cell label (ASW_label) to quantify preservation of biological structure. NMI and ARI compare annotated cell-type labels to clusters inferred from the embeddings and are computed based on *k*-means clusters. ASW_label measures within-cell-type compactness versus cross-cell-type separation in the embedding space. The aggregated bio-correction score is computed as the arithmetic mean across all three bio-correction metrics.

#### scGraph-eval metrics

The package scGraph-eval provides three metrics, Rank-PCA, Corr-PCA, and Corr-Weighted, to capture how well the embeddings preserve the cell-type relationship. Rank-PCA and Cor-PCA evaluate the Spearman and Pearson correlations with PCA-based relationships, while Corr-Weighted considers distance-based importance. In accordance with the usage guideline (<u>Usage</u>), Corr-Weighted is reported as scGraph scores. The performance of the other two metrics is reported in the Supplementary Data Table 1.

### UMAP visualization

For each embedding, we computed the neighborhood graph and generated the UMAP visualization using Scanpy v1.11.5 with identical hyperparameters. We set n_neighbors=100 and maxiter=100, while maintaining default settings for other parameters. The cells are colored by cell type annotations to compare the consistency between unsupervised clustering and known labels. For the prostate cancer dataset, the cells are also colored by assay to evaluate each method’s performance on data integration.

## Data availability

The data used in this study is not available for public distribution. To support reproducibility, a subsample of raw metrics values and source data for Fig. 1 are available with this article.

## Acknowledgement

The authors thank Weiwei Shultz (data platform) for the single cell processing pipeline and Hyejin Choi (oncology discovery) for help analyzing the prostate cancer dataset.

## Ethics declarations

This study utilizes samples that were previously collected during other research projects. In every case, informed consent was secured from participants before data collection began, and all data were anonymized to maintain confidentiality.

## Extended Data Table

**Extended Data Table 1:** Summary of benchmarking datasets.

|  | Apheresis | Prostate cancer | CAR-T drug product |
| --- | --- | --- | --- |
| data type | scRNA | scRNA, snRNA | scRNA |
| # of cells | 407,533 | 654,648 | 500,000 |
| # of sample<sup>*</sup> | 62 | 141 (57 scRNA, 84 snRNA) | 283 |
| Annotated cell types | 21 cell types: Tem/Temra cytotoxic T cells, Tem/Effector helper T cells, CD16+ NK cells, Tem/Tsen cytotoxic T cells, Non-classical monocytes, Classical monocytes, Tem/Tsen helper T cells, NKT cells, Tcm/Naive helper T cells, Regulatory T cells, MAIT, DC2, Double-positive T cells, Cycling T cells, Late erythroid, Naive B cells, Memory B cells, pDC, Tcm/Naive cytotoxic T cells, Megakaryocytes/platelets, Plasma cells | 14 cell types: Macrophages, Endothelial cells, Hepatocytes, Hepatic stellate cells, T cells, epithelial cells, Pulmonary alveolar type II cells, Smooth muscle cells, Beta cells, Luminal epithelial cells, B cells naive, Dendritic cells, Proximal tubule cells, Fibroblasts | 8 cell types: Cycling Tem/Temra cytotoxic T cells, Tem/Effector helper T cells, Cycling Tem/Effector helper T cells, Cycling Tcm/Naive cytotoxic T cells, Tcm/Naive helper T cells, Apoptotic T cells, Tcm/Naive cytotoxic T cells, Tem/Temra cytotoxic T cells |

<sup>\*</sup>We treat sample ID as the batch covariate to perform sample-level integration.

**Extended Data Table 2:** Overview of foundation models evaluated in this study.

|  | CellFM | scMulan | scFoundation | scGPT |
| --- | --- | --- | --- | --- |
| Pretraining data size | ~100 M | ~100 M | ~50 M | ~33 M |
| Model parameter size | ~800 M | ~368 M | ~100 M | ~50 M |
| Architecture | Encoder | Decoder | Asymmetric encoder-decoder | Encoder |
| Adaptation strategy | Partially fine-tuned | Zero-shot | Zero-shot | Zero-shot and fine-tuned |
| Embedding dimension | 1,536 | 1,120 | 3,072 | 512 |
| License | cc-by-nc-nd-4.0 | MIT | Code: Apache 2.0<br>Model: Proprietary | MIT |

**Extended Data Table 3:** Computation time of foundation models and baseline methods for data integration.

|  | Apheresis | Prostate cancer dataset | CAR-T drug product |
| --- | --- | --- | --- |
| Harmony | 18 mins 42 secs | 1 hour 06 mins | 34 mins 11 secs |
| scVI | 7 mins 34 secs | 7 mins 24 secs | 10 mins 01 secs |
| scGPT (zero-shot) | 1 hour 14 mins | 1 hour 37 mins | 4 hours 01 mins |
| scFoundation (zero-shot) | 17 hours 06 mins | 15 hours 48 mins | 65 hours 12 mins |
| scMulan (zero-shot) | 11 hours 23 mins | 12 hours 08 mins | 19 hours 07mins |
| scGPT (fine-tuned) | 26 hours 44 mins | 43 hours 11 mins | 32 hours 14 mins |
| CellFM (fine-tuned) | 156 hours 50 hours | 244 hours 23 min | >1000 hours* |

Computational cost was measured under a fixed evaluation environment (OS: Ubuntu 22.04.3 LTS; CPU: AMD EPYC 7R13 processor, 48 cores; RAM: 164 GiB; GPU: 4× NVIDIA L4; CUDA: 12.3). GPU was enabled where supported. Parallel runtimes adjusted to single-core equivalents.
\*Method exceeded the time limit. The reported cost is the estimate during the training process.

